# Alfalfa varieties can weakly choose beneficial nitrogen-fixing bacteria from a population isolated from a single field

**DOI:** 10.64898/2026.06.12.731664

**Authors:** Sohini Guha, Alejandra Gil-Polo, Elizabeth Paillan, Jeremy Sutherland, Ellen Bingham, Kayla M. Clouse, Liana T. Burghardt

## Abstract

1. In natural and agricultural systems, legumes recruit rhizobia from diverse soil populations to fix nitrogen in root nodules. A few legumes, including the model legume *Medicago truncatula*, can select and enrich beneficial rhizobia. Here, we investigated whether its perennial relative, *Medicago sativa* (alfalfa), a globally important forage crop, also possesses this ability.
2. We developed a genetically variable collection of 117 *Sinorhizobium meliloti* strains sampled from three field-grown alfalfa varieties, performed multi-strain and single-strain inoculations in a nitrogen-free greenhouse experiment across the same hosts, and evaluated plant benefits and relative strain fitness in nodules.
3. Alfalfa varieties differed in which strains best promoted plant growth and which strains had high fitness in nodules. Regressing strain fitness and host benefit revealed that two of three alfalfa varieties selected and enriched more beneficial strains during symbiosis, though the strength of selection was weak. In alignment with these results, no variety produced as much biomass in mixed inoculation as it did with the best-performing single strain.
4. Legumes’ ability to enrich beneficial rhizobial populations from field-representative strain diversity warrants further study to develop optimized varieties. Ultimately, identifying crop varieties that naturally select for beneficial bacteria could reduce the need for repeated inoculant applications.

## Introduction

Bio-inoculants have been used to enhance agricultural productivity for more than 120 years, with their adoption accelerating over the past 20 years (Nobbe and Hiltner, 1896; Deaker, 2004). Among the most impactful of these are nitrogen-fixing bacteria called rhizobia, which selectively form symbiotic associations with legumes (O’Callaghan et al., 2022). While external application of rhizobia to cultivated legumes can increase yields, performance can be inconsistent (Mendes et al., 2004; Mathu et al., 2012; Thilakarathna and Raizada, 2017). Roadblocks hindering the effectiveness of bio-inoculant applications include establishment and persistence in the soil and successful association with the host plant (Kaminsky et al., 2019; Mendoza-Suárez et al., 2021). The host association process can be particularly complex and is shaped by plant exudates, plant genotype, soil conditions, and even the timing of application (Venturi and Keel, 2016; Rybakova et al., 2017). Instead of striving for a single inoculant that overcomes these barriers in all environments, an alternative, underexplored approach is to utilize plant hosts that better select and reward beneficial bacteria that are already in the soil (Thies et al., 1995; Denison and Kiers, 2004; Kiers et al., 2007; Mendoza-Suárez et al., 2021; Westhoek et al., 2021). Here, we assess whether multiple varieties of alfalfa, an important forage crop, can select and reward better partners using a new rhizobial collection derived from a single agricultural field.

Legume-rhizobia symbiosis is a resource-based mutualism in which legumes provide carbon to rhizobia in exchange for nitrogen (Garau et al., 2005), thereby deriving fitness benefits from each other (Denison, 2000). The carbon rhizobia receive from plant photosynthates supports cell division and resource storage, eventually allowing rhizobia to achieve a higher cell density within nodules than in soil (Burghardt et al., 2018; Oono et al., 2020; Burghardt et al., 2026). In natural soil environments, a legume interacts with scores of genotypically diverse, co-existing rhizobia from which it chooses partners for nitrogen fixation (Gellért et al., 2026). Host-imposed selection is critical for the success of legume-rhizobia mutualism, and experiments consistently show that the fitness benefits received from the interacting partners depend on both the host and strain identity. Hosts differ in how selective they are (e.g., some hosts associate with multiple genera of rhizobia and others with a single subspecies) (Andrews and Andrews 2017). Genetic variation within a host species can also alter strain selection. This can occur both between different accessions of a single species (Burghardt et al., 2018; Mendoza-Suárez et al., 2024) and when individual genes are disrupted (Yang et al., 2010; Burghardt et al., 2020; Guha et al., 2026). Moreover, host selection of rhizobia can shift across environments. For example, nitrogen availability can shift host partner preference in *Medicago truncatula,* in which roots under greater nitrogen limitation show a stronger preference for the more efficient strains (Zhang et al., 2020). Similarly, pH shifts can dramatically change host preferences: *Sinorhizobium medicae (S. medicae)* associates with acid-soil-adapted medics while *S. meliloti* prefers medics growing in alkaline soil (Garau et al., 2005). Similarly, soybeans associate with *Sinorhizobium* in alkaline soils and *Bradyrhizobium* in acidic soils(Li et al., 2023). In sum, legume hosts vary in which rhizobia they select and host, and strain genetic variation, as well as the environment, all contribute to which strains are inside nodules.

However, just because legumes preferentially associate with certain strains does not mean that the strains they select are their most effective nitrogen-fixing partners (Porter et al., 2024). For a legume to exhibit adaptive partner choice, it must preferentially associate with or invest in certain strains over others (selectivity), and the selected strains must be beneficial ones (adaptive). Evidence of adaptive partner choice has been found for legumes like pea and *Lotus strigosus* where hosts preferentially reward a wild type fixing strain over a non-fixing mutant strain (Regus et al., 2017; Westhoek et al., 2017). But in natural and agricultural soils, strains vary continuously and subtly in symbiotic quality (Burdon et al., 1999; Gano-Cohen et al., 2019), and evidence for adaptive partner choice is less clear when hosts are given a choice among such strains. For instance, when the wild legume *Lupinus arboreus* is forced to choose between binary combinations of a good, bad, and a moderately fixing strain, originally isolated from the same host, the good strain is not consistently selected (Simms et al., 2006). Similarly, host discrimination depends on the identity of the strain population and can be relative rather than absolute—nodule occupancy of an intermediate-fixing strain shifted depending on whether it was competing against a poor fixer or a good fixer (Westhoek et al., 2021). Adaptive partner choice can also depend on host genotype (Heath and Tiffin, 2009; Bourion et al., 2018; Burghardt et al., 2018). For instance, *Medicago truncatula* accession A17 supports a more beneficial nodule community than accession R108 (Burghardt et al., 2018). Because root nodule symbiosis is context-dependent, strains that are ineffective on one genotype or environment can be beneficial in others. Taken together, the nascent picture of adaptive partner choice is not one of clear host control but of imperfect, context-dependent processes, and at present, we still have little clarity about how, when, and to what extent adaptive choice mechanisms operate within and between legume species.

Part of the reason for this gap is that it is challenging to quantify a legume’s selection and enrichment abilities in multi-strain contexts. Recent advances in genome sequencing and genome engineering have enabled the development of high-throughput methods such as Select and Resequence (SCR) and Plasmid-ID that allow the simultaneous tracking of rhizobial strain frequencies across different environments (Burghardt et al., 2018; Burghardt et al., 2020; Burghardt et al., 2022; Schumacher et al., 2025; Guha et al., 2026). The SCR technique relies on estimating strain frequencies in an initial mixed inoculum and, after selective pressure is applied, calculating relative strain fitness (log fold change). Previous selective pressures have included different host accessions, individual host mutations, and abiotic environmental conditions such as soil temperature, salinity, drought, and soil particle size (Burghardt et al., 2018; Burghardt et al., 2020; Polo et al., 2025; Guha et al., 2026). Because this high-throughput methodology simultaneously assesses scores of strains, it allows us to evaluate the host’s ability to select among strains in diverse biotic and abiotic environments. Pairing these strain-relative fitness assessments with the host-benefit data from single-strain experiments allows us to determine whether the strains that provide the highest growth benefit to the host under single-strain inoculation assays are *selected* and *enriched* when hosts have a mix of all strains to choose from (Heath and Tiffin, 2007).

Alfalfa (*Medicago sativa*) is an important legume forage crop that, due to efficient nitrogen fixation, provides nutritious forage for the dairy industry (Zhang and Wang, 2025). Alfalfa is an ideal system for assessing whether breeding for maintenance of mutualism is possible. First, it has the same host specificity as the model legume, *Medicago truncatula,* primarily associating with the same two *Sinorhizobium* species that were used to develop the SCR methodology (Young et al., 2011). Second, as a perennial, alfalfa can impose repeated selection on rhizobial populations over multiple years, unlike *Medicago truncatula,* an annual, thereby creating an opportunity to study the evolutionary consequences of a stable selection regime on rhizobia (Burghardt, 2020). The high throughput mutualism screening provided by the SCR methodology could be utilized to enable mutualism-informed breeding programs for alfalfa. If successful, this could further increase the sustainability of this key agricultural forage, especially when combined with the additional root-associated ecosystem services it provides, such as reducing nitrogen runoff and improving soil health via its deep taproot and fibrous root (Issah et al., 2020).

Here, we assessed the ability of three alfalfa varieties to choose the most beneficial partner from 117 *S. meliloti* strains that were isolated from a single alfalfa field in Pennsylvania. This strain collection is an important and novel aspect of our study—many strains in existing rhizobial collections have become lab-adapted and/or represent strains from multiple countries that may not co-occur in a single field. After isolating and sequencing this new strain collection, we performed a greenhouse experiment under nitrogen-free conditions, inoculating three alfalfa varieties, both singly and with a mix of all 117 strains. We then quantified the benefit each strain provides in single-inoculation greenhouse experiments and inferred strain frequencies in nodules on each plant to determine whether each variety selected and enriched strains that fix more nitrogen when given a choice. Specifically, we ask: 1) How genetically diverse are rhizobial strains available to hosts in a single alfalfa field at one time? 2) Do the rhizobial strains differ in the nitrogen-fixing benefits they provide to each alfalfa variety? 3) Do alfalfa varieties preferentially associate with and enrich more beneficial strains when given a choice? We found that two of three alfalfa varieties were weakly able to select and enrich better partners from the diverse strain population found in a single field.

## Materials and methods

### Development of the AVT strain collection

Sampling of alfalfa root nodules for the development of the Alfalfa Varietal Trials (AVT) strain collection occurred in late summer of 2020. We collected nodules from three alfalfa varieties (Vernal, Oneida, and SW-4107) grown at the Russell E. Larson Agricultural Research Center (Rock Springs, PA). Vernal and Onieda are both used as baseline or ‘check’ plants in the variety trials. These are varieties that are planted across many locations and years to allow performance comparisons. Vernal and Oneida were derived over 50 years ago from distinct but related breeding pools in Wisconsin and Cornell, respectively. Vernal has little pest resistance, whereas Oneida is resistant to pervasive soil pathogens (Vandemark et al., 2006). SW-4107 is a commercial variety with dormancy similar to the check varieties and high pest resistance. All varieties received Exceed Alfalfa/True Clover inoculum, and were grown in 3 × 20 ft plots arranged in a randomized block design in a field with a history of maize and alfalfa cultivation. All plots received the same phosphorus fertilizer additions over time; no extra nitrogen was added. We sampled a total of ∼100 nodules by combining ∼5-10 plants from each variety across three physically adjoining plots. We sampled nodules 4.5 years post establishment, because 1) that was when yield sampling was completed, and we were allowed to destructively sample, and 2) because at that point the rhizobial population had been exposed to the same hosts for 4 years.

Following collection, the nodules were washed with 10% bleach and rinsed three times with sterile water in a tea strainer. Next, they were ground in a sterile mortar using a pestle and 3 mL sterile phosphate-buffered saline (PBS), then filtered through a 10-um filter into a clean tube. The filter was rinsed with PBS, and 400 μL of the filter wash was added to each tube. The nodule homogenate was then serially diluted in Yeast-Mannitol (YM) broth or Reasoner’s 2A Broth (R2A) broth to obtain a cell density of 10^9^ CFU/mL. This was further diluted 1:10,000 and mixed with resazurin (Prospector® Dye Reagent), which was added to a final concentration of 100 µM. Finally, the samples were vacuum loaded onto a Prosector® array, which was sealed according to the manufacturer’s instructions. After incubating the cultures at three time points – 0, 1, and 5 days, Prospector personnel transferred 51% of the samples to a larger-volume destination plate, where isolate-level growth yielded 2627 isolates. We then subcultured 371 wells, focusing on the 5-day time point. We purified them three times on YM plates, then grew the isolated colonies in Tryptone Yeast (TY) broth and stored the liquid cultures in 25% glycerol at -80°C. To narrow the collection down to the final 117 strains that were the most proteomically diverse, isolates were subjected to Matrix-assisted laser desorption/ionization time-of-flight (MALDI-TOF) protein mass spectrometry and the strains with distinct protein spectra were selected. Lastly, we used 16S rRNA Sanger sequencing (Forward primer_27F: 5’ AGAGTTTGATCCTGGCTCAG 3’, Reverse primer 342R : 5’ CTGCTGCSYCCCGTAG 3’) to confirm these strains belonged to the genus *Sinorhizobium*.

### Long-read sequencing and genome assembly of the strain collection

We sequenced all 117 strains on the PacBio Sequel IIe sequencing platform (Pacific Biosciences) at the Penn State Huck Institutes of Life Sciences Genomic Core Facility, which produces long, highly accurate DNA reads (HiFi reads). Sequencing generated ∼48,000–57,000 reads per strain (3.3–4.0 × 10⁸ bases), corresponding to 50-60X coverage per isolate, assuming a genome size of ∼6.7 Mb. Assembly was performed with hifiasm v0.25.0 using default parameters to generate primary and alternate contig sets (Cheng et al., 2021). Genome completeness, contamination, and assembly quality metrics (including N50; median 3.93 Mb, range 2.01–5.60 Mb), contig count (median 4, range 2–6), genome size (mean 7.45 Mb, range 5.43–8.03 Mb), and GC content (mean 62.0%) were assessed with CheckM2 v1.1.0 (Chklovski et al., 2023). All assemblies exhibited very high completeness (median 100%, range 98.9–100%) and low contamination (median 0.75%, range 0–3.04%). Given these high-quality metrics, no additional pre-annotation filtering for contaminating contigs was performed. Based on genome assembly quality and comparative coverage across the replicons, AVT118 was chosen as the in-house reference genome for downstream analyses.

### Gene prediction, genome annotation, and average nucleotide identity

Gene prediction and functional annotation of the assemblies were performed using Prokka v1.14.6 with its default Prodigal v2.6.3 gene caller (Hyatt et al., 2010; Seemann, 2014). Pairwise average nucleotide identity (ANI) was calculated across 117 strain assemblies using the ANIm method as implemented in pyANI package v0.2.12 (Pritchard et al., 2016). Genome nucleotide sequences were extracted from the GFF files prior to analysis. All pairwise ANI comparisons were performed using NUCmer from the MUMmer 3.0 package, with default parameters using shell script run_anim_from_gff.sh (Marçais et al., 2018).

### Pangenome construction for comparative genomics

To evaluate the core and accessory gene contents of the strains, annotated assemblies were analyzed with Panaroo v1.5.2 to construct a pan-genome (Tonkin-Hill et al., 2020). Panaroo was run with strict cleaning of spurious annotations (--clean-mode strict), MAFFT as the aligner (--aligner mafft), and generation of a core-gene alignment (--alignment core).

### Defining the core-soft core-shell-cloud genes categories and calculating the presence-absence variance

The Core (>99%); Soft Core (99%-95%); Shell (95%-15%); Cloud (< 15%) genes were defined based on (Gonzalez-Diaz et al., 2022). The Presence-Absence variance (PAV) matrix was evaluated based on the accessory (Soft Core+Shell+Cloud) genes. In-house custom Python scripts (pangenome_accessory_figure.py and pangenome_ucurve.py) were used to map the PAV matrix against the AVT strain phylogeny and calculate the distribution of the accessory gene categories and individual accessory genes within the strain collection.

### Replicon assignment of the AVT strains and PAV distribution on the replicons

Replicon assignment to the AVT strain genomes was performed using a custom shell pipeline composed of three shell scripts: 1) code1_deduplicate_reference_db.sh: removes redundant sequences from a *S. meliloti* reference genome database composed of 139 strains downloaded from NCBI, 2) code2_homogenize_via_rm1021.sh: standardizes replicon nomenclature by BLASTing the deduplicated reference sequences against *S. meliloti* RM1021, ensuring uniform nomenclature of the plasmids in the database, and 3) code3_classify_avt_strains.sh: parses assembly contigs to record the contig number and length. Contigs were then queried via BLASTn against the reference database. Contigs <100,000 bp were flagged as assembly fragments and merged into their corresponding replicon category based on percent identity (Fragment_); all remaining unassigned contigs were classified as accessory plasmids (Plasmid_other). Following the replicon assignment, the PAV data were superimposed on the newly annotated replicons to determine the PAV proportions on the replicons.

### Single copy core gene identification

A gene was classified as core if it was present (non-empty, non-missing entry) in all 117 AVT strains. Genes for which any strain carried a semicolon-delimited entry-indicating multiple paralogs detected by Panaroo were additionally excluded. To identify genes suitable for concatenated phylogenetic inference, the same single-copy core gene were applied to the combined set of 117 AVT strains and two *S. meliloti* reference genomes (RM1021 and USDA1106).

### Maximum likelihood-based phylogenetic tree construction

Single-copy core genes were extracted and concatenated using custom R v4.6.1 scripts from the Panaroo PAV output for all AVT genomes and the two additional reference genomes, which served as the outgroups.

Briefly, for each gene in the alignment directory, we retained only those alignment files in which all target strains were represented exactly once. Alignments passing these filters were concatenated in order, with sequences from each strain appended end-to-end to produce a single multi-locus alignment. The concatenated alignment was used as input to construct the maximum-likelihood phylogenetic tree using IQ-TREE 3 (Wong et al., 2025). The best-fit nucleotide substitution model was determined using ModelFinder (–m MFP), which evaluates multiple candidate models and selects the optimal model based on Bayesian Information Criterion (Kalyaanamoorthy et al., 2017). The selected model (MK+FQ+I+R9) was used for tree reconstruction for this study. Branch support was assessed using two complementary approaches: ultrafast bootstrap approximation (UFBoot2) with 1000 replicates (–bb 1000) and Shimodaira-Hasegawa (SH)-like approximate likelihood ratio test (SH-aLRT) with 1000 replicates (–alrt 1000).

### Variant calling on the AVT genomes

To call variants based on the reference genome, we wrote a custom script to convert the long reads to 250 bp short reads that retained quality information for downstream analyses. Specifically, we combined long reads with spacer characters and created 250-bp chunks until a spacer was reached. This resulted in a FASTQ file for each strain. We then mapped these short reads to our reference genome AVT118 to call variants using bwa mem v0.7.17 (Li & Durbin, 2009) with 20 threads. Read group information was assigned during mapping using the -R flag with sample identifiers. Resulting alignments were coordinate sorted with samtools sort (Li et al., 2009) and were indexed with samtools index. Variants were called jointly across all 117 strains using bcftools mpileup followed by bcftools call (v1.23.1) (Danecek et al., 2021) with the --ploidy 1 flag to correctly model the haploid bacterial genome. Only variant sites were retained (-mv flag). Pairwise SNP differences between all strain pairs were calculated using bcftools gtcheck on genotype calls converted to biallelic format following normalization with bcftools norm -m -.

### Inoculum preparation for the single and multi-strain inoculation experiments in the greenhouse

We performed a two-pronged experiment at the same time in a greenhouse at Penn State. For the single-strain inoculation experiment, we grew each AVT strain separately for 48 hours in 2 mL Tryptone-Yeast extract (TY) media at 28°C, shaking at 120 rpm. We adjusted the optical density of each strain culture to 0.1 (OD_600_). The inoculum was prepared by diluting 4 mL of the OD_600_-adjusted culture in 50 mL of DI water. A total of 4 mL was applied to each pot, eight days after seed sowing. Each pot received approximately 10^7^ CFU. For the multi-strain inoculation experiment, we mixed 100 μL of each strain’s OD_600_-adjusted culture, brought the volume to 100 mL with sterile DI water, and inoculated each pot with 4 mL of this solution (∼10^7^ CFU per pot). A subset of the inoculum was reserved for genomic DNA isolation, which was performed by centrifuging eight 1 mL aliquots at 18,000 x g for 2 min, removing the supernatant, and freezing the pellet at -20°C prior to DNA extraction. Initial communities were sequenced, and frequencies were inferred as outlined in the Rhizobial Fitness section below.

### Alfalfa germination and maintenance conditions in the greenhouse

For the single-strain inoculation experiment, we grew four replicates of each of the three alfalfa varieties (Vernal, Oneida, and SW-4107) in nitrogen-limited conditions (N = 1140; 4 reps x 3 varieties x 117 isolates) (Fig. 1). For the multi-strain inoculation experiment, we planted 6 replicates per alfalfa variety (N = 18; 6 reps x 3 varieties). Single-strain and multi-strain experiments were conducted in the same greenhouse bay but on separate benches. Before planting, seeds were surface sterilized by soaking in a 10% bleach solution for 90 seconds, then rinsed in deionized water.

**Figure 1:**
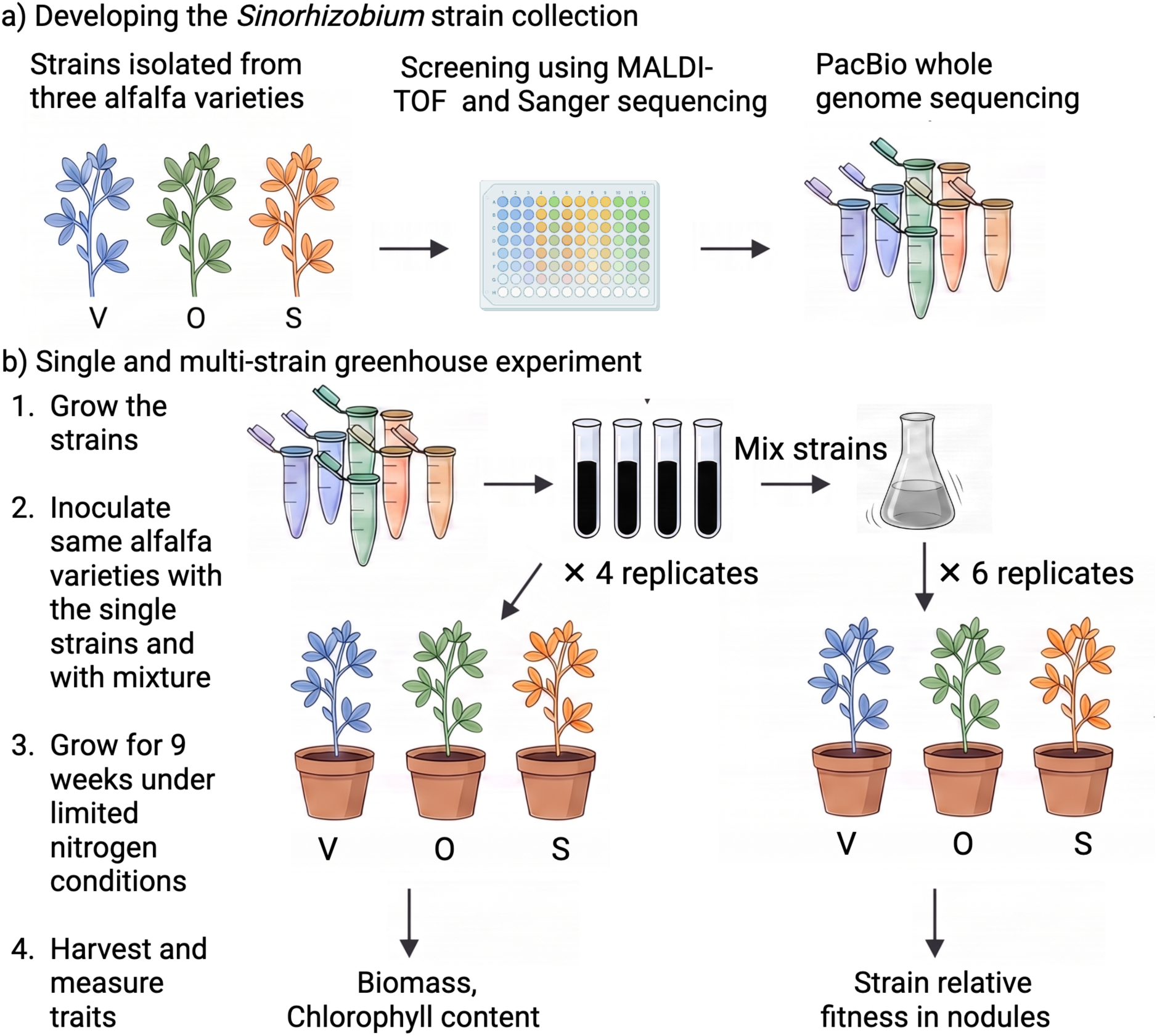
Overview of strain collection development and greenhouse experiment to evaluate host benefits and rhizobial success. (A) *Development of the AVT strain collection* - *Sinorhizobium meliloti* strains were isolated from nodules in an alfalfa variety trial at Penn State University. Pools of ∼100 nodules were sampled from 5-10 alfalfa plants of each of the following varieties (Oneida (O), Vernal (V), and the commercial variety SW-4107 (S)). We used the Prospector system to obtain over 2,000 putative isolates. We selected 117 strains from the larger collection based on unique protein profiles as indicated by a Matrix-assisted laser desorption/ionization time-of-flight (MALDI-TOF) protein spectra analysis and confirmed membership in the genus *Sinorhizobium* using Sanger sequencing of 16S rRNA. Lastly, strains were sequenced using long-read PacBio technology and assembled using hifiasm v0.25.0 using default parameters. (B) *Greenhouse experiment*- We grew cultures of each of the 117 strains and used them to conduct two complementary randomized block experiments in nitrogen-free conditions. First, a single-strain experiment in which four replicate pots of each of the three host varieties were inoculated with each strain individually. 9 weeks post-inoculation, we measured chlorophyll content, above-ground biomass, below-ground biomass, and nodule count. Second, a mixed inoculation experiment in which all 117 strains were combined in equal amounts. Six replicate pots of each of the three varieties were inoculated with the mixture, and 9 weeks post-inoculation, we measured the same traits. In addition, all nodules were dissected from the roots, rhizobia were enriched from the pools, rhizobial genomic DNA was extracted (Burghardt et al., 2018), and the pools were sequenced using Illumina short-read sequencing. The resulting reads were mapped to an in-house reference genome (AVT 118) to determine relative strain frequencies and evaluate individual strain fitness across the three varieties. Created in https://BioRender.com

Surface-sterilized seeds were stratified on wet filter paper at 4°C in the dark for 2 days, then planted in duplicate in Deepot™ Support Tray (D20T for 2.5” & 2.7”; item number: D20T), filled with a 50:50 sand-calcined clay media, arranged in four blocks for the single-strain and six for the multi-strain experiment. The seeds were covered with ¼ to ½ inch of sterile vermiculite. The pots were kept moist with twice-daily misting, and humidity was maintained during germination by covering them with a clear plastic tarp. After one week, pots were moved to their randomized locations and inoculated. Each pot was watered as needed, with care taken to avoid cross-contamination between pots. The pots were fertilized weekly with 2x Fahräeus medium (David G. Barker et al., 2006). Uninoculated plants were included in each block to monitor cross-contamination from watering. Four weeks post-sowing, we thinned all pots to one plant per pot. Nine weeks post-inoculation, we harvested the plants.

### Measurement of plant traits

Chlorophyll content and total biomass were selected as proxies to evaluate plant performance. These traits were measured in both the single- and mixed-inoculation experiments. The chlorophyll content for each replicate was measured using an MC-100 Chlorophyll Concentration Meter (Apogee Instruments) 9 weeks post-inoculation. We measured the chlorophyll content of three leaves per plant and averaged the values. Total biomass/plant was measured by weighing the above-ground (shoot) and the below-ground (root) biomass together after oven-drying at 75 °C for 3 days.

### Assessment of relative strain fitness

Rhizobial relative fitness was evaluated only for the multi-strain inoculation experiment. To assess rhizobial relative fitness, we pooled all nodules from individual alfalfa plants, surface-sterilized them in 10% bleach for 1 min and then rinsed them thoroughly in DI water. Nodule pools were then homogenized for 90 s in 4 mL of sterile 0.85% sodium chloride solution using an Omni TH tissue homogenizer. The nodule homogenate was then spun at 400 x g for 10 min at 10 °C to pellet the bacteroids and plant debris. We collected the supernatant enriched in undifferentiated rhizobial cells, pelleted them at 18,000 x g for 5 min, pipetted off the supernatant, and froze the pellets at -20 °C until DNA extraction. DNA was extracted using a Qiagen DNeasy® Plant extraction kit following the manufacturer’s protocol, except we used 3 μL of RNase in the first step and 50 μL of nuclease water for the final elution. Samples were sequenced using Illumina short-read technology at the Penn State Huck Institutes of Life Sciences Genomic Core Facility. Indexed libraries were generated using the Illumina DNA Prep Kit. Libraries were pooled and sequenced on a 150 x 150 paired-end NextSeq 1000 High Output run, resulting in an average read depth of ∼75. We trimmed the reads using TrimGalore! v0.4.1 (www.bioinformatics.babraham.ac.uk/projects/trim_galore/) with default settings and minimum read length: 100 bp. Quality threshold was 30 (−-quality); and minimum adapter match was 3 (−stringency). We used bwa mem v0.7.17 with default settings to align reads to AVT118, and strain frequencies were estimated using HARP. Strain frequencies were used to calculate the *relative strain fitness* of each strain by taking the log_2_ of the ratio of the final strain frequency in the nodules to the initial frequency in the inoculum (median initial frequency = 0.0090, 5%-95% quantile: 0.006–0.01). The logarithmic transformation puts proportional increases and decreases on the same scale (e.g., a fourfold decrease and increase have values of −6 and 6, respectively). Strains with frequency estimates of 0 in the selected communities were assigned a fitness value of −6, less than the lowest measurable reductions in other strains.

### Statistical analyses

All analyses were run in the R statistical environment v4.3.2. We used linear models with analysis of variance (ANOVA) to test the effects of host variety, strain identity, and their interaction on total biomass per plant (Total Biomass∼ Strain*Variety) and chlorophyll content per plant (Chlorophyll Content∼ Strain*Variety) as the proxies for host benefit. For all the linear models, replicates were included as an additive predictor. To monitor strain selection across varieties, we performed an RDA using the rda() function from the vegan v2.6-4 package, with relative strain fitness as the response matrix and variety as the explanatory variable. The significance of each model was assessed by extracting the adjusted *R*^2^ and via permutation-based ANOVA (1,000 permutations) using the anova() function with by = “terms”. Predictors with *P* value < 0.05 indicated significant changes in relative strain fitness. To identify which host variety pairs drove the overall signal, we performed pairwise RDAs for all three variety combinations (O vs V, O vs S, V vs S) using the same approach, with the host variety as the sole explanatory variable in each subset. Significance was assessed via permutation-based ANOVA (9999 permutations), and adjusted *R*^2^ (Adj.*R*^2^) and proportion of variance explained were extracted for each comparison.

## Results

### High genetic variability among the AVT strains is observed at the gene and SNP levels

To characterize the breadth of genetic variation in this unique strain collection, we assessed gene content variation, phylogenetic relatedness, and single-nucleotide polymorphisms.

Whole-genome sequencing and Average Nucleotide Identity (ANI) analysis confirmed that all 117 strains wer*e S. meliloti* with ANI values >99% (except strains AVT18 and AVT 105) (Fig S1, Table S1). We detected approximately 12,135 gene clusters across the entire pangenome, which can be categorized by their occurrence in the strain collection, of which 3,925 were present in all strains, 1,809 in 95–99% of strains, 2,464 in 15–95% of strains, and 3,937 in <15% of strains (Figure 2, Table S2). *S. meliloti* success in the soil and rhizosphere (diCenzo et al., 2014). As expected, genes on the chromosome had the highest likelihood of being present in all strains, at 48.4%. Interestingly, while 39.9% of pSymB genes were found in all strains, only a few pSymA genes (0.1%) were shared across all strains (Figure S2). pSymA harbored the highest proportion of Shell genes, while the Cloud genes were found across all replicons but were most prevalent on accessory plasmids as expected (Figure S2, Table S2). Examination of genetic diversity at the SNP level revealed 57,076 bi-allelic SNPs on the chromosome, 34,349 on pSymA, and 48,722 on pSymB (Table S3). Of the replicons, the chromosome had the lowest SNP densities (14.58 SNPs/kb, e.g., was most conserved) while pSymB was the most polymorphic (29.44 SNPs/kb) (Table S4). The minimum number of SNPs differentiating two strains was 16 SNPs (AVT49 vs AVT48), whereas the maximum was 76,504 SNPS (AVT18 vs AVT78) (Table S5). Interestingly, we found that strains from all three host varieties are dispersed across a maximum-likelihood phylogeny based on SNP variation in 3774 single-copy core genes and across presence-absence variation (Figure 2). In sum, despite being collected at a single time point from a single field, this collection is genetically diverse at both the gene-content and sequence levels, making it an interesting resource for assessing adaptive host choice.

**Figure 2:**
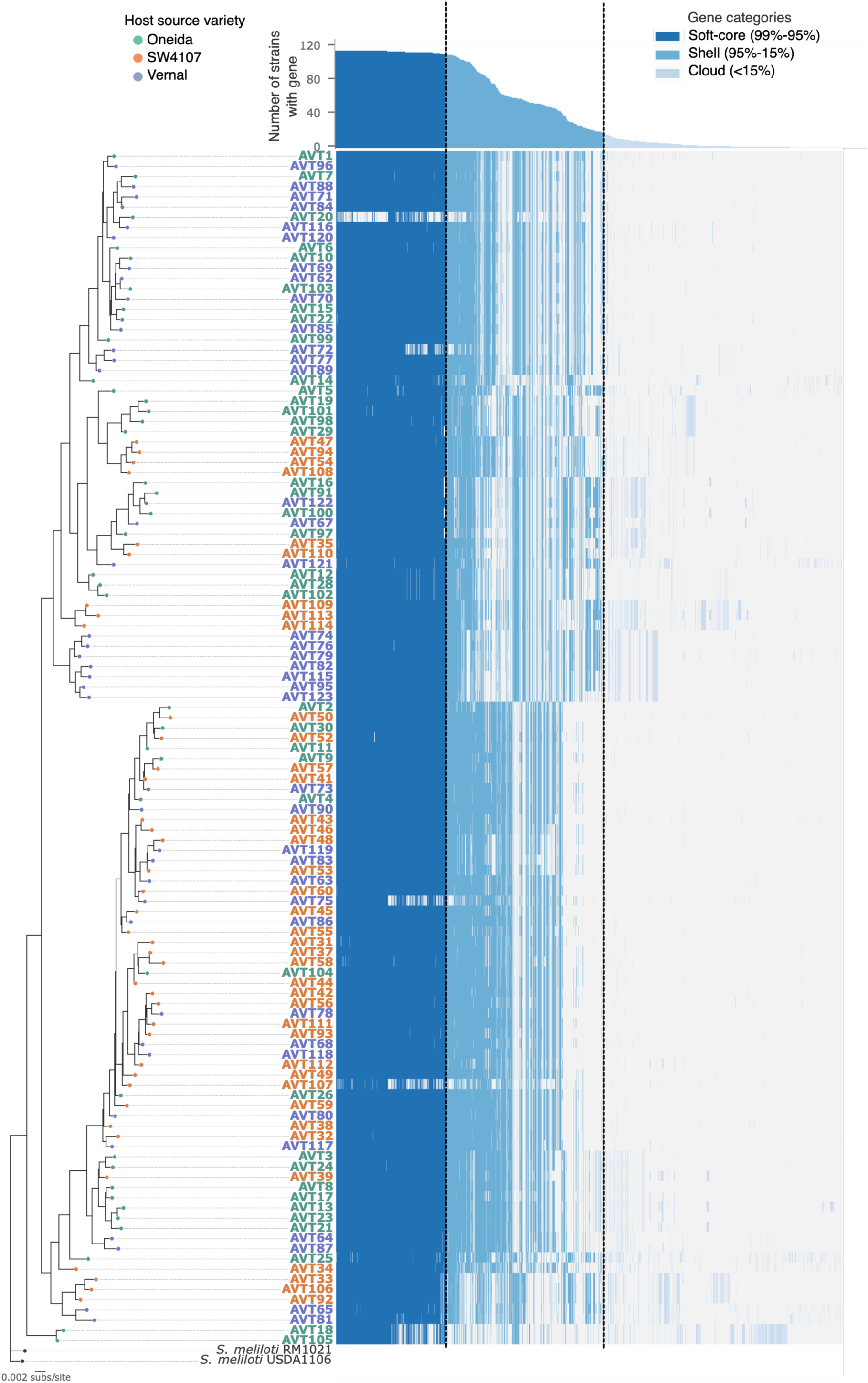
Phylogenetic relatedness and gene content variation in 117 strains sampled from a single alfalfa field: After using Panaroo to determine the pangenome of the strains, we generated a maximum likelihood tree from 3774 single-copy core genes using IQ-TREE 3.0.1, which showed three deep subclades. Popular strains USDA1106 and RM1021 were included as reference and were used as the outgroups to root the tree. The branch tip colours denote the original variety of alfalfa from which the strain was isolated [orange=SW4107(S), green= Oneida (O), purple = Vernal (V)]. The best fit substitution model (MK+FQ+I+R9) was selected based on the Bayesian Information Criterion. Branch support of the tree was assessed using two independent metrics: ultrafast bootstrap (UFBoot2; 1000 replicates) and the SH-like approximate likelihood ratio test (SH-aLRT; 1000 replicates). Accessory gene presence or absence on the right side of the figure revealed differences in the gene content across the isolates. The PAV matrix is based on ∼5000 accessory genes, which include the Soft Core (99%-95%), Shell (95-15%), and Cloud (< 15%) (Gonzalez-Diaz et al., 2022).

### Strains vary widely in the benefits provided to hosts, and beneficial strains vary by host

Variation in nitrogen fixation efficiency was assessed using host biomass acquisition and chlorophyll content in nitrogen-free conditions, two commonly used proxies for symbiotic performance (Fritschi and Ray, 2007; Heath and Tiffin, 2009). For host biomass acquisition, strain, host, and their interaction explained 49% of trait variation. The significant interaction between strain identity and host variety (*P* value = 0.02) indicates that a given strain has different effects across host varieties (Fig 3 A-C). Chlorophyll content, an indicator of nitrogen content in leaves, showed a broadly similar pattern for strain-level effects (27% of the variance, *P* value <0.001) and revealed a significant main effect of host variety (6% of the variance, *P* value <0.001) (Fig S4). However, unlike biomass, no significant strain-by-variety interaction was detected for chlorophyll content (*P* value = 0.53) (Table 1). Taken together, our results suggest that the pool of strains alfalfa hosts have to choose from in a single alfalfa field varies widely in their nitrogen-fixing abilities, and the strains that are most beneficial tend to differ by host.

**Figure 3:**
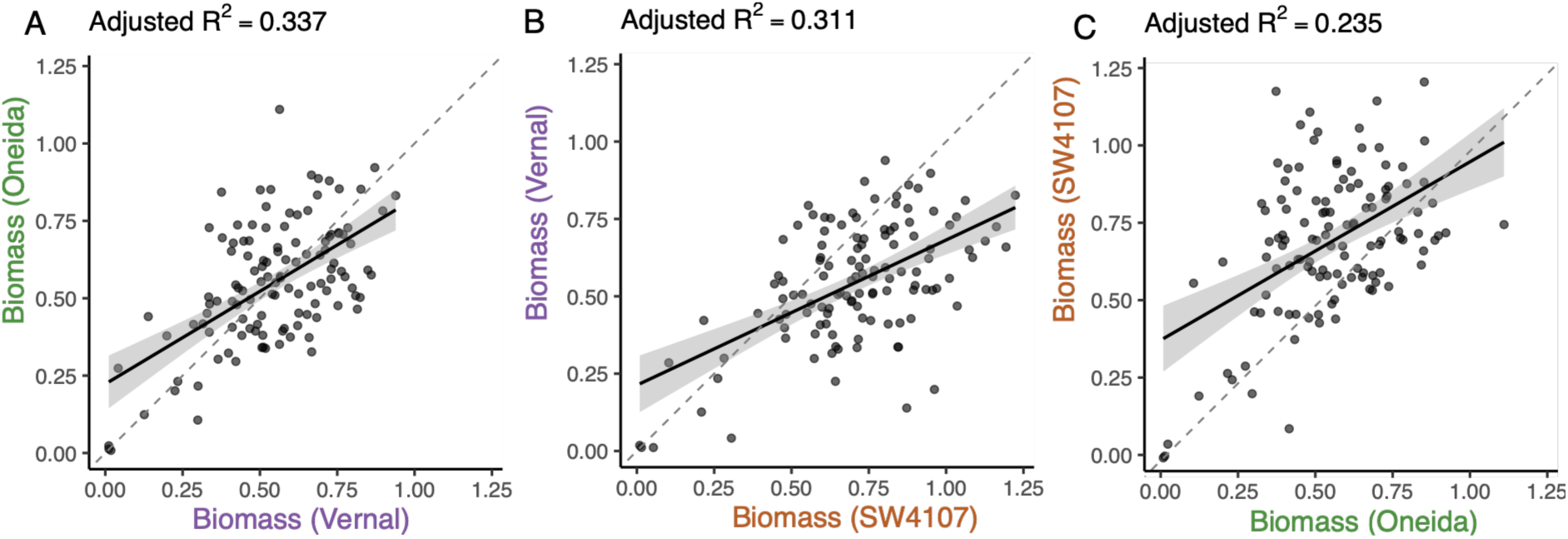
AVT strains vary in host benefits conferred: Scatter plots showing the relationship between the growth benefits (biomass) each strain provides to each host variety in nitrogen-free conditions (A-C). Each point reflects the mean plant biomass across 4 replicate pots. The dotted line represents y = x, and the solid line is the regression line; the shaded area indicates the 95% confidence interval. See Table 1 for model results

**Table 1:** ANOVA results for linear models of host benefits from the AVT strains.

| Model and terms | Df<sup>2</sup> | %Var<sup>3</sup> | F value<sup>4</sup> | P value<sup>5</sup> |
| --- | --- | --- | --- | --- |
| *Biomass/plant*(Adj.R<sup>2</sup>=0.34)<sup>1 ***</sup> |  |  |  |  |
| Strain | 116 | 28.1 | 5.7 | < 0.001*** |
| Variety | 2 | 6 | 54.5 | < 0.001*** |
| Rep | 3 | 4 | 21.7 | < 0.001*** |
| Strain × Variety | 232 | 15 | 1.2 | 0.02* |
| Residuals | 911 | 47.7 | NA | NA |
| *Chlorophyll/plant* (adj.R<sup>2</sup>=0.28)<sup>***</sup> |  |  |  |  |
| Strain | 116 | 26.7 | 5.2 | < 0.001*** |
| Variety | 2 | 6 | 55.5 | < 0.001*** |
| Rep | 3 | 2 | 11.9 | < 0.001*** |
| Strain × Variety | 232 | 13.2 | 0.9 | 0.5 |
| Residuals | 911 | 51 | NA | NA |

<sup>1</sup>Adj. R<sup>2</sup>, adjusted coefficient of determination for the full model;
<sup>2</sup>Df= degrees of freedom;
<sup>3</sup>%Var = Percentage of total variance explained by each predictor;
<sup>4</sup>Fval= F-statistic from the ANOVA
<sup>5</sup>PVAL= Significance level of the predictor effect.
Significance codes: \*\*\*P < 0.001; \*\*P < 0.01; \*P < 0.05; .P < 0.1; ns, not significant.
NA= not applicable.

### Alfalfa varieties differ in their ability to associate with the most beneficial strains when given a choice

To determine which rhizobial strains hosts select and enrich when given a choice, we performed a multi-strain inoculation experiment. Each host variety was inoculated with a mixture of all 117 strains, and we quantified the relative fitness of each strain within the pooled nodules using the Select-and-Resequence approach (Burghardt et al., 2018). A redundancy analysis, constrained by host variety and replicate (Fig 3D, Fig S6, Table S6), followed by an ANOVA-like permutation-based test, revealed that host variety alone explained 36.1% of the variation in strain fitness (*P* value = 0.01, full model adj R² = 0.11, Table S7). Pairwise comparisons further revealed that strain fitness differed significantly between Oneida and Vernal (Adj R² = 0.19, *P* value = 0.002) and between Vernal and the commercial variety (Adj R² = 0.12, *P* value = 0.01) but not between Oneida and the commercial variety (O vs S: Adj R² ≈ 0, *P* value = 0.37) (Table S8). However, we note that there is variability across pots, even within a host variety, in which strains show high (or low) relative fitness in a host’s nodules (Fig S7). This pattern could be due to genetic variation within a single alfalfa variety or to unaccounted environmental variation (Fig S7).

To determine whether host varieties could select and enrich beneficial strains in their nodules, we ran linear models regressing relative strain fitness from the multi-strain inoculation experiment on host-benefit data from the single-strain inoculation experiment (Fig 4). For the Oneida variety, strain fitness in the mixed inoculations was positively correlated with host benefit (Adj R² = 0.12, *P* = 0.0001). While weaker, the commercial variety also showed a significant positive correlation (Adj R² = 0.04, *P* = 0.01). In contrast, Vernal exhibited no association between strain fitness and host benefit (Adj R² = 0.01, *P* = 0.1). Because of our observation of high pot replicate-level variation in strain relative fitness, we also calculated regression lines for each pot. We observe similar correlation trends for each variety at the pot level. Although we observed a negative relationship between strain fitness and host benefits, Vernal recruited a compositionally distinct strain community(Fig 4D). This decoupling suggests that Vernal is unable to select for strains based on mutualistic quality within a diverse field mixture. Taken together, we show that some alfalfa varieties can preferentially select and enrich beneficial strains from a realistic pool of naturally varying field strains, but this ability is weak and varies across hosts.

**Figure 4:**
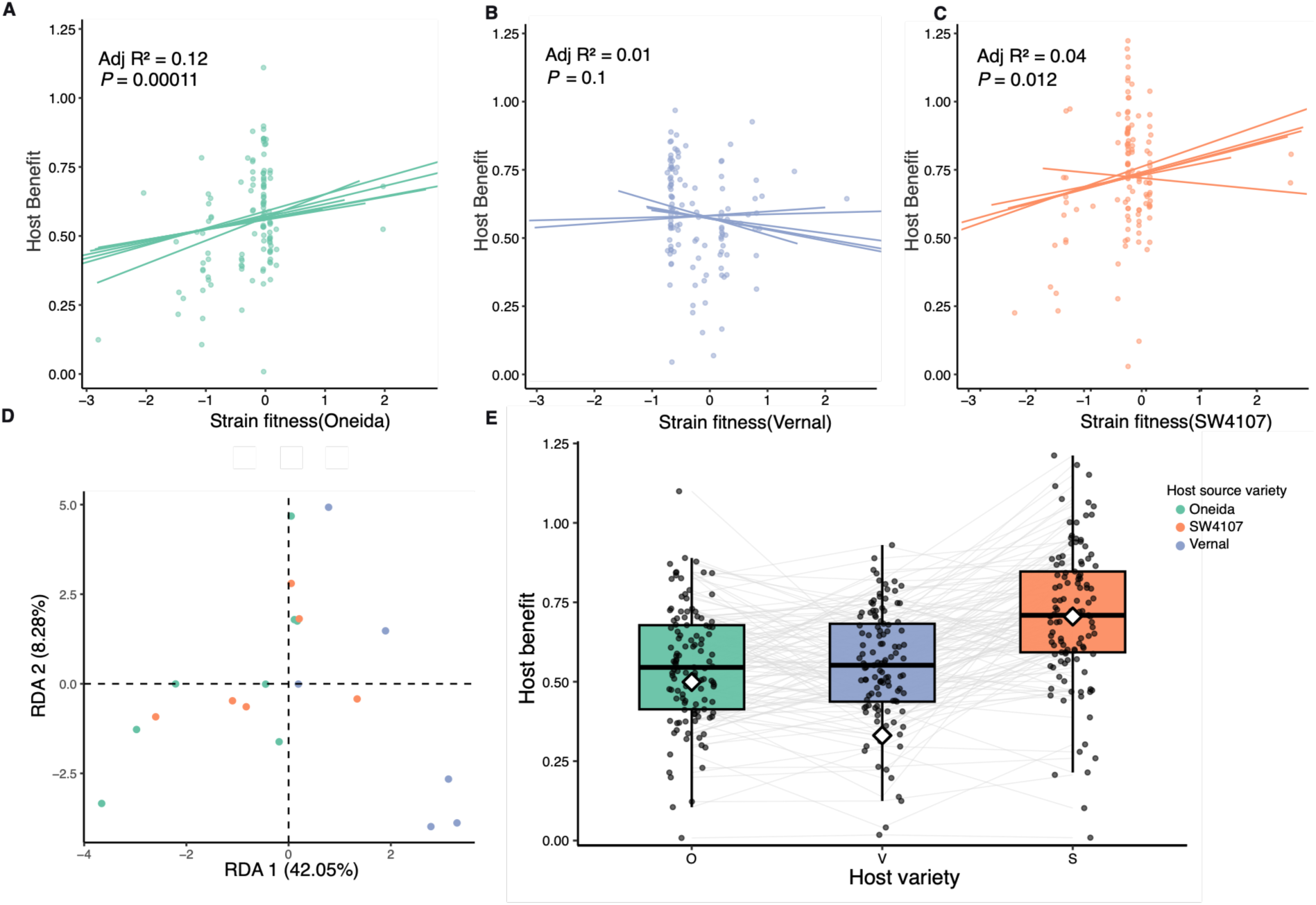
Alfalfa varieties differ in their ability to select beneficial partners from a mixture. (A-C) Relationship between strain relative fitness during multi-strain inoculations and host benefit provided by each strain as inferred from host biomass in nitrogen-free single-strain inoculation experiments. Points represent mean strain relative fitness and mean host benefits summarized across replicates to provide the overall pattern, while lines represent the estimated relationship for each replicate pot separately to visualize the relationship consistency (or not) across replicates. For model details, refer to Table S7. (D) Host identity explains ∼50.33% of the total variation in strain relative fitness. Here we show the first two axes from a Redundancy Analysis (relative strain fitness ∼ host variety). Points denote replicate pots, and colours are coded by host variety. Refer to Fig. S7 for details on the relative fitness of individual strains across replicates across host varieties. (E) No host varieties were able to derive as much benefit in a mixture (à = mean biomass acquired during multi-strain inoculation) as they did with the best single strain (·= mean biomass acquired during single strain inoculation with each strain), but Vernal, the host that did not exhibit the ability to select and enrich beneficial strains in (B), was particularly growth-limited.

## Discussion

This study addresses a foundational challenge in sustainable agriculture: whether legume varieties can preferentially enrich beneficial symbionts and whether it is possible to maximise nitrogen fixation in field contexts by selecting host varieties better equipped to choose partners with higher nitrogen fixation abilities. Using an ecologically and agronomically important crop, alfalfa (*Medicago sativa*), and its bacterial symbiont, *S. meliloti*, we show that alfalfa’s capacity to enrich for beneficial rhizobial strains from a diverse rhizobial community differs across varieties. Our results reveal that 1) even in a single field, hosts have a genetically diverse population of strain partners to choose from both in terms of gene presence-absence and SNP diversity 2) these strains vary substantially in the benefits they provide to hosts, with the most beneficial strains differing between alfalfa varieties 3) two of three alfalfa varieties were able to select and enrich more beneficial strains during symbiosis, though the strength of selection is weak and variable. These findings highlight the nuanced interplay between legume host varieties and their symbionts and underscore the importance of considering both partners in efforts to optimize nitrogen fixation. By integrating host-mediated microbial selection into plant breeding strategies, we can better harness the full potential of plant–microbe mutualisms for sustainable agriculture.

### High rhizobial diversity within a small field site

Our strain collection, which was sampled from a small area at a single time, has a diversity level consistent with the high genomic variation observed in past studies of strain collections sampled across continents (Epstein et al., 2012; Epstein and Tiffin, 2021). Similarly, at a comparatively narrower geographic scale, a collection of 191 strains of *S. meliloti* sampled from 21 sites across the native Mediterranean range of *Medicago truncatula* demonstrated high local genomic variation and minimal isolation-by-distance, indicating that even geographically constrained populations harbor considerable standing diversity (Riley et al., 2021). Earlier studies evaluating natural genetic variation within *S. meliloti* populations from single sites have been limited either by small sample sizes or by the use of Multi Locus Sequence Typing (MLST) rather than whole-genome sequencing. Despite these limitations, (Van Berkum et al., 2007) detected as many as 21 sequence types through MLST in a collection of 148 isolates from a single site. In contrast, (Toro et al., 2017)sampled 21 *S. meliloti* from a single site, but found them to be largely clonal. Here, we expand upon their work by using a larger sample and high-quality genome assemblies to demonstrate a high level of intraspecific genetic diversity within a single *S. meliloti* population. The observed intraspecific diversity, while striking, is not limited to *S. meliloti* and has been previously observed within *Rhizobium leguminosarum* and *Bradyrhizobium* sp., sampled from spatially constrained sites (Kumar et al., 2015; Weisberg et al., 2022). While the AVT collection showed a high degree of overall nucleotide similarity (ANI > 99%), SNP variation and gene presence-absence data captured natural genetic variation occurring in nodules located within 3 meters of each other in a single field. A similar pattern has been documented across other members of the *Rhizobiales,* where comparative genomics across multiple species has shown that core genomes comprise only 10% of the genome, with accessory genes being an important source of adaptive variation (Galardini et al., 2011; Rosselli et al., 2021). The documented genetic diversity reflects the complexity of the soil population each host chooses from, as well as the competition any introduced strain must face (Kaminsky et al., 2019). Understanding how hosts cope with and interact with this diversity requires continued characterization of diversity at the individual field level (Thies et al., 1995; Burghardt, 2020).

### Nitrogen fixation outcomes are strongly affected by standing natural genetic variation within a rhizobial population and among host varieties

Our study showed a significant strain-by-variety interaction, reaffirming the importance of genotype-by-genotype interactions in shaping symbiosis outcomes (Heath and Tiffin, 2009; Burghardt et al., 2018; Burghardt et al., 2020; Guha et al., 2026). The substantial variation observed in individual strain performance across host varieties further reaffirms the importance of strain-level variation in determining symbiotic outcomes in *Medicago*. In the *Sinorhizobium*-*Medicago* system, strain-specific effects on nitrogen fixation have been previously documented, ranging from complete loss to subtle quantitative variation in effectiveness (Heath and Tiffin, 2007; Rangin et al., 2008; Heath and Tiffin, 2009). Moreover, early cross-inoculation studies of multiple *Medicago* genotypes demonstrated that nitrogen-fixation effectiveness depended on the *S. meliloti* or *S. medicae* strains they associated with (Heath and Tiffin, 2007). The wide variation we found in nitrogen fixation outcomes of the AVT strains aligns with previous observations of host-specific strain performance variation and significant strain-by-host interactions between *S. meliloti* strains and hosts from the same sampling site (Heath and Tiffin, 2009). In this study, we go a step further to find functional variation in nitrogen fixation at a very fine spatial scale: nodules collected at a single time point, sampled within 3 meters of each other, and find that most rhizobial lineages were represented in nodules of all three host varieties. These observations point to a broader agronomic challenge: a strain that performs well in one variety may be mismatched in another (Wendlandt et al., 2019). This has direct implications for the use of rhizobial inoculants in agriculture, where a few elite strains selected for high nitrogen-fixation abilities are broadly deployed across genetically diverse crops. While striking, the diversity of strain-by-host outcomes we observe is perhaps unsurprising, as our host varieties had co-occurred and selected on this strain population for at least 4.5 years, and these Pennsylvania alfalfa fields have a long-term history of alfalfa production, as well as the presence of agricultural weeds such as *Melilotus alba* and *Medicago lupulina,* which also associate with *Sinorhizobium*.

### Alfalfa varieties differ in their ability to choose beneficial partners from a field strain community

Whether and how legume hosts selectively associate with more beneficial rhizobial partners, i.e., do they exhibit adaptive partner choice, has long been a central question in the legume– rhizobia literature, with mixed empirical support that varies across host species, rhizobial strains, and environmental contexts (Porter et al., 2024). Here, we find that two out of the three tested alfalfa varieties were able, albeit weakly, to select and enrich beneficial strains from a diverse, field-derived strain community. Our results for alfalfa align with several previous studies of adaptive partner choice in other *Medicago* species. For instance, four *M. truncatula* lines inoculated with 225 *Sinorhizobium* isolates from a single Mediterranean soil demonstrated that nodule communities were influenced by plant genotype and bacterial competition for nodulation in addition to the fixation efficiencies of the bacterial genotypes (Rangin et al., 2008). Building on previous results showing that three different *Medicago* genotypes were weakly able to select and enrich beneficial strains from complex communities (Burghardt et al., 2018; Batstone et al., 2020), Burghardt et al. (2026) found that one mechanism underlying the adaptive partner-choice trait was that larger nodules tended to harbor more beneficial strains, and *Medicago* genotypes differed in this ability.. Another approach to infer the presence of adaptive partner choice is to compare plant biomass when plants are given a choice with that when they are given the most effective strains. Results in *Medicago* also vary based on this metric. While Rangin et al. (2008) observed that inoculation with the *S. medicae* community across multiple *Medicago* genotypes produced plants with biomass levels comparable to those of the most efficient strain in the absence of competition. Heath and Tiffin (2009) observed that *Medicago truncatula* plants were smaller when inoculated with a community of *S. meliloti* than with each strain individually. In our study, none of the alfalfa varieties derived as much benefit from the mixture as they did from the best single strains, suggesting ample opportunity to further improve adaptive partner choice. To our knowledge, our study provides the first test of adaptive partner choice against an ecologically realistic strain pool, and our observation that two of the three can enrich beneficial strains, although weakly, suggesting that this trait is variable and potentially breedable in an agronomically important species.

## Conclusions

Our finding of adaptive partner choice in alfalfa, a forage crop relative of the model species *Medicago truncatula*, strengthens the observation that adaptive partner choice is a common, if imperfect, feature of *Medicago–Sinorhizobium* symbiosis. Expanded screening of existing commercial varieties against multiple locally sourced rhizobial communities can help identify varieties with superior adaptive partner-choice capacity. In the longer term, quantifying the heritability of host benefits and partner-choice traits in alfalfa using QTL mapping and GWAS approaches analogous to those already applied in soybean, common bean, and *Medicago truncatula* could enable identification of genomic regions for marker-assisted selection for adaptive partner choice (Gorton et al., 2012; Kamfwa et al., 2019; Torkamaneh et al., 2020; Batstone et al., 2022). Such methods could accelerate the integration of rhizobial choice as a breeding target alongside yield, disease resistance, and stress tolerance (Li and Brummer, 2012; Mendoza-Suárez et al., 2024). Moving forward, breeding for host partner choice abilities could be paired with the introduction of variety-specific elite inoculum development to create sustainable agro-evolutionary systems. While domestication has reshaped plant genomes, it has also reshaped microbiomes, raising the possibility of breeding strategies extending to microbiomes (De Zutter and Audenaert, 2026). Encouragingly, ours and others’ data suggest breeders may be working with the evolutionary grain rather than against it—recent evidence suggests that legume and rhizobium fitness interests are largely aligned (Batstone et al., 2022; Epstein et al., 2023). If adaptive partner choice is indeed a tractable, heritable trait in forage legumes, it could ultimately serve as a lever for reducing nitrogen fertilizer dependency while simultaneously ensuring the stability of mutualism.

## Acknowledgments

We thank Patrick Sydow, Amanda Jason, Andy Swartley, and Jennifer E. Harris for their assistance with the tedious task of picking and processing nodules, Russel E Larson Agricultural Station for hosting the sourced field locations, Scott DiLoreto, greenhouse manager for greenhouse assistance, Tyler Rice for supporting sampling of strains from the alfalfa variety trials, Genomics Core Facility of Huck Institutes of the Life Sciences (RRID: 365 SCR_023645) at Pennsylvania State University for performing the whole genome sequencing of the strain collections, the high performance research computing cluster (ROAR-COLLAB) managed by Institute for Computational and Data Sciences (ICDS) at the Pennsylvania State University for providing support for performing all the high throughput data analysis.

## Authorship Credits

LTB idea conception and funding, and experimental design, EB, ELP, LTB, MAG, SG, plant care, data collection for main experiment; JS, SG performed mapping and variant calling, KMC assembled the genomes, ran the Panaroo analyses, generated the Presence-Absence matrix; SG performed the Harp analysis, replicon assigning, phylogeny, and statistical tests; SG and LTB wrote the first draft, followed by extensive feedback from MAG and KMC; all authors reviewed and edited the manuscript.

## Data Availability

All raw sequencing reads from the initial rhizobial strain community and from rhizobial genomic DNA extracted from the nodule pools of the individual host varieties have been deposited in NCBI under Bioproject PRJNA1482181 (Submission ID: SUB16286289). All strain assemblies will be deposited in NCBI. All raw data and analysis code have been uploaded to GitHub which and will be made public (https://github.com/sohini5981/AVT_SS) upon publication. The source code and raw data will be available in a public repository (e.g., Dryad) upon publication as well.

## References

Batstone RT, Burghardt LT, Heath KD (2022) Phenotypic and genomic signatures of interspecies cooperation and conflict in naturally occurring isolates of a model plant symbiont. Proc R Soc B Biol Sci 289: 20220477

Batstone RT, O’Brien AM, Harrison TL, Frederickson ME (2020) Experimental evolution makes microbes more cooperative with their local host genotype. Science 370: 476–478

Bourion V, Heulin-Gotty K, Aubert V, Tisseyre P, Chabert-Martinello M, Pervent M, Delaitre C, Vile D, Siol M, Duc G, et al (2018) Co-inoculation of a Pea Core-Collection with Diverse Rhizobial Strains Shows Competitiveness for Nodulation and Efficiency of Nitrogen Fixation Are Distinct traits in the Interaction. Front Plant Sci 8: 2249

Burdon JJ, Gibson AH, Searle SD, Woods MJ, Brockwell J (1999) Variation in the effectiveness of symbiotic associations between native rhizobia and temperate Australian Acacia: within-species interactions. J Appl Ecol 36: 398–408

Burghardt LT (2020) Evolving together, evolving apart: measuring the fitness of rhizobial bacteria in and out of symbiosis with leguminous plants. New Phytol 228: 28–34

Burghardt LT, Epstein B, Guhlin J, Nelson MS, Taylor MR, Young ND, Sadowsky MJ, Tiffin P (2018) Select and resequence reveals relative fitness of bacteria in symbiotic and free-living environments. Proc Natl Acad Sci 115: 2425–2430

Burghardt LT, Epstein B, Hoge M, Trujillo DI, Tiffin P (2022) Host-Associated Rhizobial Fitness: Dependence on Nitrogen, Density, Community Complexity, and Legume Genotype. Appl Environ Microbiol 88: e00526–22

Burghardt LT, Sydow P, Sutherland J, Epstein B, Tiffin P (2026) Genetic variation in host selectivity and adaptive strain enrichment in legume-rhizobia symbiosis: host-dependent, imperfect processes correlate with nodule morphology. Proc R Soc B Biol Sci 293: 20252851

Burghardt LT, Trujillo DI, Epstein B, Tiffin P, Young ND (2020) A Select and Resequence Approach Reveals Strain-Specific Effects of *Medicago* Nodule-Specific PLAT-Domain Genes. Plant Physiol 182: 463–471

Cheng H, Concepcion GT, Feng X, Zhang H, Li H (2021) Haplotype-resolved de novo assembly using phased assembly graphs with hifiasm. Nat Methods 18: 170–175

Chklovski A, Parks DH, Woodcroft BJ, Tyson GW (2023) CheckM2: a rapid, scalable and accurate tool for assessing microbial genome quality using machine learning. Nat Methods 20: 1203–1212

Danecek P, Bonfield JK, Liddle J, Marshall J, Ohan V, Pollard MO, Whitwham A, Keane T, McCarthy SA, Davies RM, et al (2021) Twelve years of SAMtools and BCFtools. GigaScience 10: giab008

Deaker R (2004) Legume seed inoculation technology?a review. Soil Biol Biochem 36: 1275–1288

Denison RF (2000) Legume sanctions and the evolution of symbiotic cooperation by rhizobia. Am Nat 156: 567–576

diCenzo GC, MacLean AM, Milunovic B, Golding GB, Finan TM (2014) Examination of prokaryotic multipartite genome evolution through experimental genome reduction. PLOS Genet 10: e1004742

Epstein B, Branca A, Mudge J, Bharti AK, Briskine R, Farmer AD, Sugawara M, Young ND, Sadowsky MJ, Tiffin P (2012) Population Genomics of the Facultatively Mutualistic Bacteria Sinorhizobium meliloti and S. medicae. PLoS Genet 8: e1002868

Epstein B, Tiffin P (2021) Comparative genomics reveals high rates of horizontal transfer and strong purifying selection on rhizobial symbiosis genes. Proc R Soc B Biol Sci 288: 20201804

Fritschi FB, Ray JD (2007) Soybean leaf nitrogen, chlorophyll content, and chlorophyll a/b ratio. Photosynthetica 45: 92–98

Galardini M, Mengoni A, Brilli M, Pini F, Fioravanti A, Lucas S, Lapidus A, Cheng J-F, Goodwin L, Pitluck S, et al (2011) Exploring the symbiotic pangenome of the nitrogen-fixing bacterium Sinorhizobium meliloti. BMC Genomics 12: 235

Gano-Cohen KA, Wendlandt CE, Stokes PJ, Blanton MA, Quides KW, Zomorrodian A, Adinata ES, Sachs JL (2019) Interspecific conflict and the evolution of ineffective rhizobia. Ecol Lett 22: 914–924

Garau G, Reeve WG, Brau L, Deiana P, Yates RJ, James D, Tiwari R, O’Hara GW, Howieson JG (2005) The Symbiotic Requirements of Different Medicago Spp. Suggest the Evolution of Sinorhizobium Meliloti and S. Medicae with Hosts Differentially Adapted to Soil pH. Plant Soil 276: 263–277

Gellért C, Ebrahimkhalili N, Siwakoti S, Zhu H, Kereszt A (2026) Not All Allies Are Welcome: Partner Discrimination in Legume–Rhizobium Symbiosis. Mol Plant-Microbe Interactions® MPMI-08–25-0108-FI

Gonzalez-Diaz A, Carrera-Salinas A, Pinto M, Cubero M, Van Der Ende A, Langereis JD, Domínguez MÁ, Ardanuy C, Bajanca-Lavado P, Marti S (2022) Comparative pangenome analysis of capsulated Haemophilus influenzae serotype f highlights their high genomic stability. Sci Rep 12: 3189

Gorton AJ, Heath KD, Pilet-Nayel M-L, Baranger A, Stinchcombe JR (2012) Mapping the Genetic Basis of Symbiotic Variation in Legume-Rhizobium Interactions in *Medicago truncatula*. G3 GenesGenomesGenetics 2: 1291–1303

Guha S, Bledsoe RB, Sutherland J, Epstein B, Fry GM, Venugopal V, Sankari S, Gil-Polo A, Levin G, Geddes BA, et al (2026) Mutations in legume genes that influence symbiosis create a complex selective landscape for rhizobial symbionts. ISME J 20: wrag005

Heath KD, Tiffin P (2009) Stabilizing mechanisms in a legume-rhizobium mutualism. Evolution 63: 652–662

Heath KD, Tiffin P (2007) Context dependence in the coevolution of plant and rhizobial mutualists. Proc R Soc B Biol Sci 274: 1905–1912

Hyatt D, Chen G-L, LoCascio PF, Land ML, Larimer FW, Hauser LJ (2010) Prodigal: prokaryotic gene recognition and translation initiation site identification. BMC Bioinformatics 11: 119

Issah G, Schoenau JJ, Lardner HA, Knight JD (2020) Nitrogen Fixation and Resource Partitioning in Alfalfa (Medicago sativa L.), Cicer Milkvetch (Astragalus cicer L.) and Sainfoin (Onobrychis viciifolia Scop.) Using 15N Enrichment under Controlled Environment Conditions. Agronomy 10: 1438

Kalyaanamoorthy S, Minh BQ, Wong TKF, Von Haeseler A, Jermiin LS (2017) ModelFinder: fast model selection for accurate phylogenetic estimates. Nat Methods 14: 587–589

Kamfwa K, Cichy KA, Kelly JD (2019) Identification of quantitative trait loci for symbiotic nitrogen fixation in common bean. Theor Appl Genet 132: 1375–1387

Kaminsky LM, Trexler RV, Malik RJ, Hockett KL, Bell TH (2019) The inherent conflicts in developing soil microbial inoculants. Trends Biotechnol 37: 140–151

Kessner D, Turner TL, Novembre J (2013) Maximum likelihood estimation of frequencies of known haplotypes from pooled sequence data. Mol Biol Evol 30: 1145– 1158

Kiers ET, Hutton MG, Denison RF (2007) Human selection and the relaxation of legume defences against ineffective rhizobia. Proc R Soc B Biol Sci 274: 3119–3126

Kumar N, Lad G, Giuntini E, Kaye ME, Udomwong P, Shamsani NJ, Young JPW, Bailly X (2015) Bacterial genospecies that are not ecologically coherent: population genomics of *Rhizobium leguminosarum*. Open Biol 5: 140133

Li X, Brummer EC (2012) Applied Genetics and Genomics in Alfalfa Breeding. Agronomy 2: 40–61

Li Y, Wang C, Zheng L, Ma W, Li M, Guo Z, Zhao Q, Zhang K, Liu R, Liu Y, et al (2023) Natural variation of GmRj2/Rfg1 determines symbiont differentiation in soybean. Curr Biol 33: 2478–2490.e5

Marçais G, Delcher AL, Phillippy AM, Coston R, Salzberg SL, Zimin A (2018) MUMmer4: A fast and versatile genome alignment system. PLOS Comput Biol 14: e1005944

Mathu S, Herrmann L, Pypers P, Matiru V, Mwirichia R, Lesueur D (2012) Potential of indigenous bradyrhizobia versus commercial inoculants to improve cowpea (*Vigna unguiculata* L. walp.) and green gram (Vigna radiata L. wilczek.) yields in Kenya. Soil Sci Plant Nutr 58: 750–763

Mendes IC, Hungria M, Vargas MAT (2004) Establishment of Bradyrhizobium japonicum and B. elkanii strains in a Brazilian Cerrado oxisol. Biol Fertil Soils 40: 28–35

Mendoza-Suárez M, Akyol TY, Nadzieja M, Andersen SU (2024) Increased diversity of beneficial rhizobia enhances faba bean growth. Nat Commun 15: 10673

Mendoza-Suárez M, Andersen SU, Poole PS, Sánchez-Cañizares C (2021) Competition, nodule occupancy, and persistence of inoculant strains: key factors in the *Rhizobium*-legume symbioses. Front Plant Sci 12: 690567

Nobbe F, Hiltner L (1896) Inoculation of the Soil for Cultivating. US Patent 570 813.

O’Callaghan M, Ballard RA, Wright D (2022) Soil microbial inoculants for sustainable agriculture: Limitations and opportunities. Soil Use Manag 38: 1340–1369

Oono R, Muller KE, Ho R, Jimenez Salinas A, Denison RF (2020) How do less-expensive nitrogen alternatives affect legume sanctions on rhizobia? Ecol Evol 10: 10645–10656

Polo MAG, Bledsoe RB, Calvert MB, Cherry L, Epstein B, Fudge R, Harris J, Tiffin P, Burghardt LT (2025) Rhizobia independently adapt to soil and legume host environments, but soil conditions influence the abundance of high-quality partners. 2025.12.08.692960

Porter SS, Dupin SE, Denison RF, Kiers ET, Sachs JL (2024) Host-imposed control mechanisms in legume–rhizobia symbiosis. Nat Microbiol 9: 1929–1939

Pritchard L, Glover RH, Humphris S, Elphinstone JG, Toth IK (2016) Genomics and taxonomy in diagnostics for food security: soft-rotting enterobacterial plant pathogens. Anal Methods 8: 12–24

Rangin C, Brunel B, Cleyet-Marel J-C, Perrineau M-M, Béna G (2008) Effects of *medicago truncatula* genetic diversity, rhizobial competition, and strain effectiveness on the diversity of a natural *sinorhizobium* species community. Appl Environ Microbiol 74: 5653–5661

Riley AB, Grillo MA, Epstein B, Tiffin P, Heath KD (2021) Partners in space: Discordant population structure between legume hosts and rhizobium symbionts in their native range. doi: 10.1101/2021.06.30.449460

Rosselli R, La Porta N, Muresu R, Stevanato P, Concheri G, Squartini A (2021) Pangenomics of the Symbiotic Rhizobiales. Core and Accessory Functions Across a Group Endowed with High Levels of Genomic Plasticity. Microorganisms 9: 407

Rybakova D, Mancinelli R, Wikström M, Birch-Jensen A-S, Postma J, Ehlers R-U, Goertz S, Berg G (2017) The structure of the Brassica napus seed microbiome is cultivar-dependent and affects the interactions of symbionts and pathogens. Microbiome 5: 104

Schumacher JD, Dusek N, Mendoza-Suárez M, Geddes BA (2025) Adaptation of plasmid-ID technology for evaluation of N2-fixing effectiveness and competitiveness for root nodulation in the sinorhizobium–medicago system. Environ Microbiol 27: e70118

Seemann T (2014) Prokka: rapid prokaryotic genome annotation. Bioinformatics 30: 2068–2069

Simms EL, Taylor DL, Povich J, Shefferson RP, Sachs JL, Urbina M, Tausczik Y (2006) An empirical test of partner choice mechanisms in a wild legume–rhizobium interaction. Proc R Soc B Biol Sci 273: 77–81

Thies JE, Woomer PL, Singleton PW (1995) Enrichment of *Bradyrhizobium* spp populations in soil due to cropping of the homologous host legume. Soil Biol Biochem 27: 633–636

Thilakarathna MS, Raizada MN (2017) A meta-analysis of the effectiveness of diverse rhizobia inoculants on soybean traits under field conditions. Soil Biol Biochem 105: 177–196

Tonkin-Hill G, MacAlasdair N, Ruis C, Weimann A, Horesh G, Lees JA, Gladstone RA, Lo S, Beaudoin C, Floto RA, et al (2020) Producing polished prokaryotic pangenomes with the Panaroo pipeline. Genome Biol 21: 180

Torkamaneh D, Chalifour F-P, Beauchamp CJ, Agrama H, Boahen S, Maaroufi H, Rajcan I, Belzile F (2020) Genome-wide association analyses reveal the genetic basis of biomass accumulation under symbiotic nitrogen fixation in African soybean. Theor Appl Genet 133: 665–676

Toro N, Villadas PJ, Molina-Sánchez MD, Navarro-Gómez P, Vinardell JM, Cuesta-Berrio L, Rodríguez-Carvajal MA (2017) The underlying process of early ecological and genetic differentiation in a facultative mutualistic Sinorhizobium meliloti population. Sci Rep 7: 675

Van Berkum P, Badri Y, Elia P, Aouani ME, Eardly BD (2007) Chromosomal and Symbiotic Relationships of Rhizobia Nodulating *Medicago truncatula* and *M. laciniata*. Appl Environ Microbiol 73: 7597–7604

Vandemark GJ, Ariss JJ, Bauchan GA, Larsen RC, Hughes TJ (2006) Estimating genetic relationships among historical sources of alfalfa germplasm and selected cultivars with sequence related amplified polymorphisms. Euphytica 152: 9–16

Venturi V, Keel C (2016) Signaling in the Rhizosphere. Trends Plant Sci 21: 187–198

Weisberg AJ, Rahman A, Backus D, Tyavanagimatt P, Chang JH, Sachs JL (2022) Pangenome Evolution Reconciles Robustness and Instability of Rhizobial Symbiosis. mBio 13: e00074–22

Wendlandt CE, Regus JU, Gano-Cohen KA, Hollowell AC, Quides KW, Lyu JY, Adinata ES, Sachs JL (2019) Host investment into symbiosis varies among genotypes of the legume Acmispon strigosus, but host sanctions are uniform. New Phytol 221: 446– 458

Westhoek A, Clark LJ, Culbert M, Dalchau N, Griffiths M, Jorrin B, Karunakaran R, Ledermann R, Tkacz A, Webb I, et al (2021) Conditional sanctioning in a legume– *Rhizobium* mutualism. Proc Natl Acad Sci 118: e2025760118

Wong T, Ly-Trong N, Ren H, Baños H, Roger A, Susko E, Bielow C, De Maio N, Goldman N, Hahn M, et al (2025) IQ-TREE 3: Phylogenomic Inference Software using Complex Evolutionary Models. doi: 10.32942/X2P62N

Yang S, Tang F, Gao M, Krishnan HB, Zhu H (2010) *R* gene-controlled host specificity in the legume–rhizobia symbiosis. Proc Natl Acad Sci 107: 18735–18740

Zhang Y, Wang L (2025) Advances in basic biology of alfalfa (*Medicago sativa L.*): a comprehensive overview. Hortic Res 12: uhaf081

