## Supplemental Methods and Figs for "Alfalfa varieties can weakly choose beneficial nitrogen-fixing bacteria from a population isolated from a single field"

Running Title: Breeding for mutualisms

Keywords: *Medicago sativa*, relative strain fitness, adaptive partner choice, rhizobia, legume, microbial, mutualism, genomes, *Sinorhizobium meliloti*

ORCID

E. Paillan 0000-0001-7159-4721
A. Gil-Polo 0000-0001-8715-5176
S. Guha 0000-0002-6914-1093
L. Burghardt 0000-0002-0239-1071
K. Clouse 0000-0003-0047-8920
J. Sutherland 0000-0003-3535-0302

Supplementary Figures

Figure S1: Pairwise ANI values among all AVT strains
Figure S2: Distribution of the Core, Soft core, Shell and Cloud genes within the AVT strain collections
Figure S3: Frequency of occurrence of the core, soft-core, shell and cloud genes within the AVT strains.
Figure S4**:** AVT strains exhibiting variations in conferring host benefits (chlorophyll)
Figure S5: Histogram showing the distribution of the frequencies of 117 *S. meliloti* strains in the initial community used to inoculate the three alfalfa hosts.
Figure S6**:** Redundancy Analysis (relative strain fitness ~ host) showing the third and fourth RDA axes
Figure S7: Heat map of relative strain fitness (strain log2-fold change) across the individual nodule pools from each plant (6 plants per variety), ordered according to the maximum-likelihood based core gene phylogeny of the AVT strains.

Supplemental Tables

Table S1- The average nucleotide identity values of the strains (including the outgroups) included in the study (provided as a .tsv file).
Table S2- Distribution of the core, softcore, shell, cloud genes within the AVT strains across the three replicons
Table S3- SNP (biallelic) counts across the replicons of the AVT strains based on pangenome
Table S4- Normalised SNP counts on the replicons
Table S5- Pairwise SNP mismatches (provided as a .csv file).
Table S6- Raw frequency counts/ relative strain fitness (provided as a .tsv file)
Table S7- RDA paired with PERMANOVA analysis of nodule isolate fitness across the hosts
Table S8-Pairwise RDA comparison of relative strain fitness across the varieties

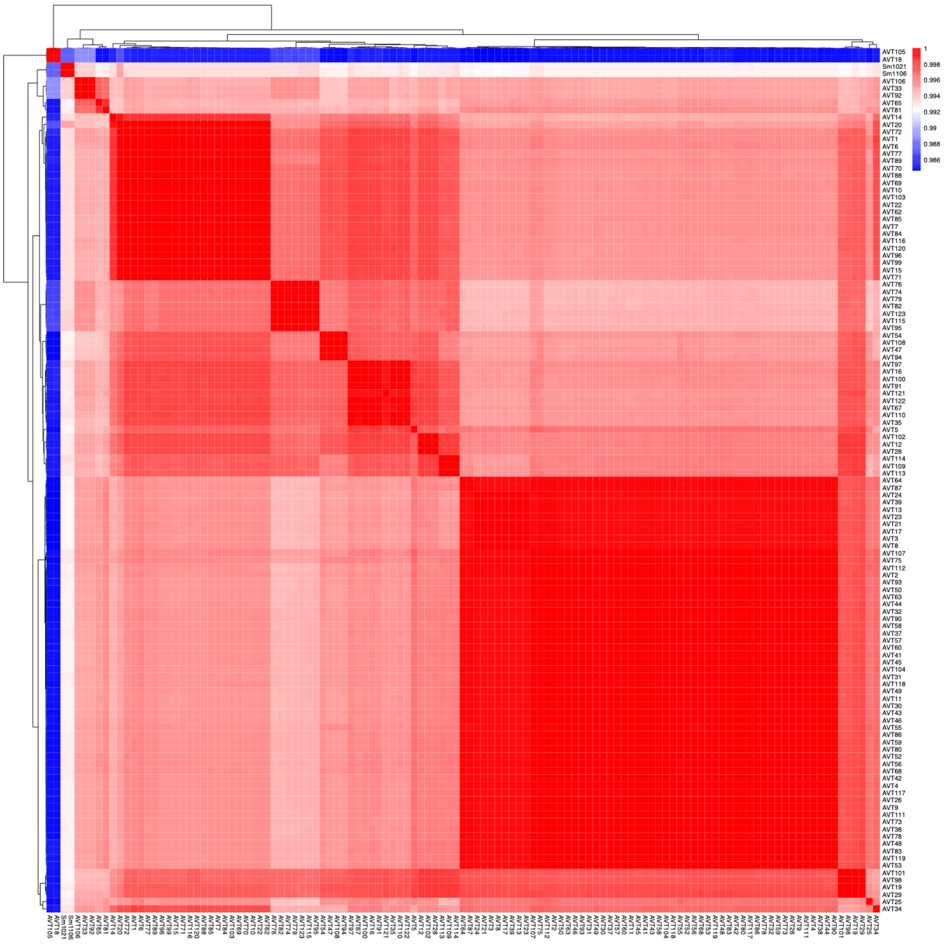

**Figure S1:** Pairwise ANI values among all AVT strains were calculated from whole-genome comparisons and visualized as a hierarchical clustering heatmap. Each cell represents the ANI between two strains, with darker red indicating higher nucleotide identity and lighter shades indicating lower identity. Rows and columns are ordered according to hierarchical clustering based on pairwise ANI distances. For details, refer to Table S1

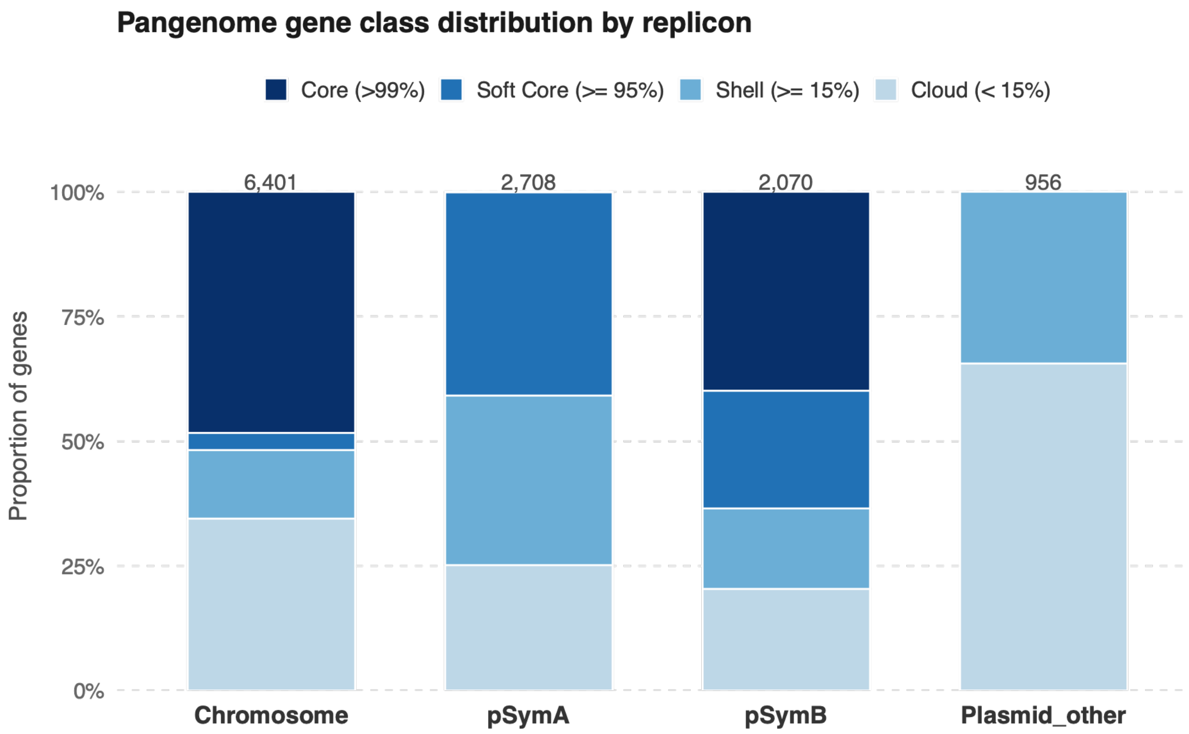


**Figure S2:** Distribution of the Core, Soft core, Shell, and Cloud genes within the AVT strain collections across the replicons. Each gene category has been defined according to the definition outlined in (Gonzalez-Diaz et al., 2022) and is color-coded according to the key provided. The numbers at the top of each bar plot indicate the total number of genes present on each replicon. For more details, see Table S2.

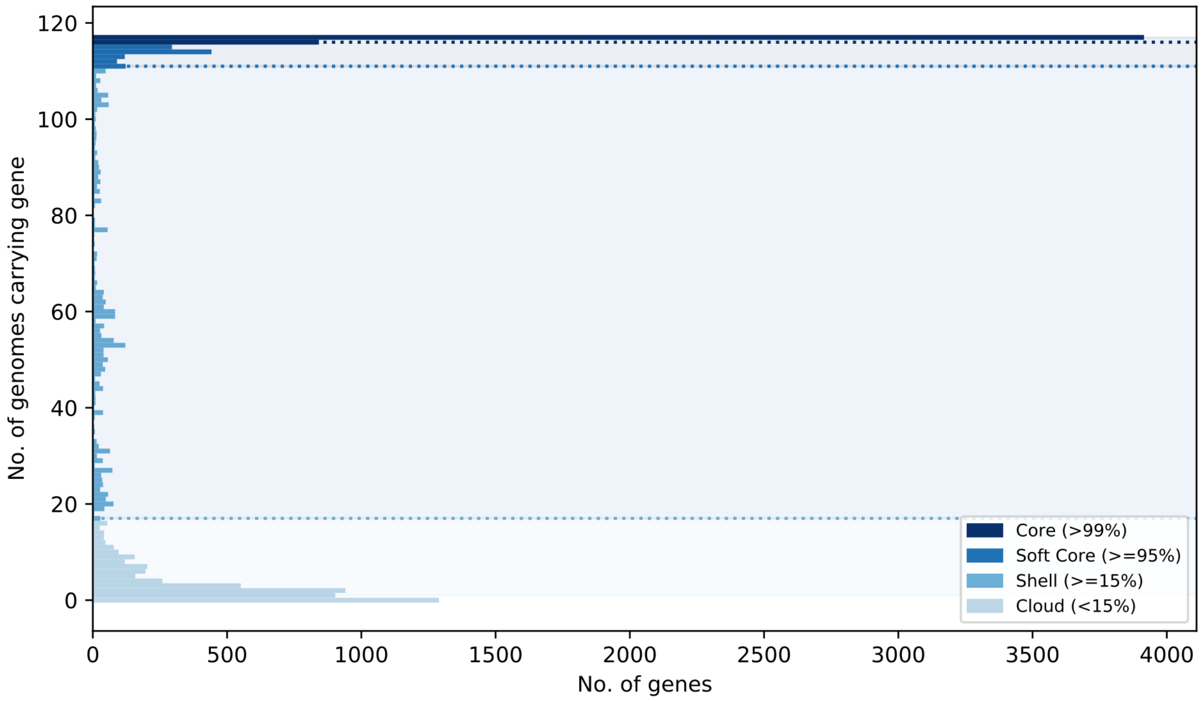


**Figure S3:** Frequency of occurrence of the core, soft-core, shell, and cloud genes within the AVT strains. The x-axis indicates the total number of genes, and the y-axis indicates the number of genomes carrying genes belonging to each category. Bars are coloured by gene category: dark blue indicates core genes (>99% of genomes), medium blue indicates soft core genes (≥95% of genomes), light blue indicates shell genes (≥15% of genomes), and pale blue indicates cloud genes (<15% of genomes).

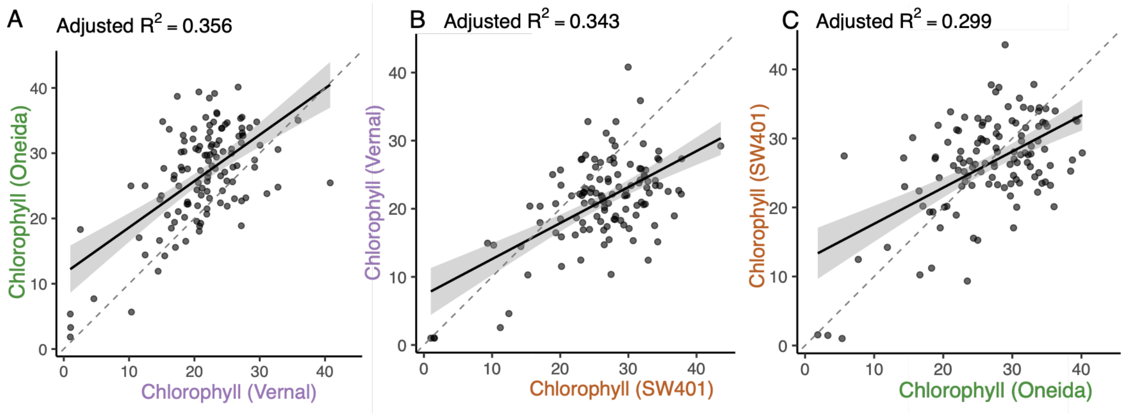


**Figure S4:** AVT strains exhibiting variations in conferring host benefits (chlorophyll): Scatter plots showing correlations between strain performances based on the mean chlorophyll content per strain in each of the three host varieties (A-C). The dotted line indicates y=x, and the solid line represents the line of regression, with the shaded area indicating the 95% confidence interval. All three models were statistically significant. For model details, refer to Table 1 and S6.

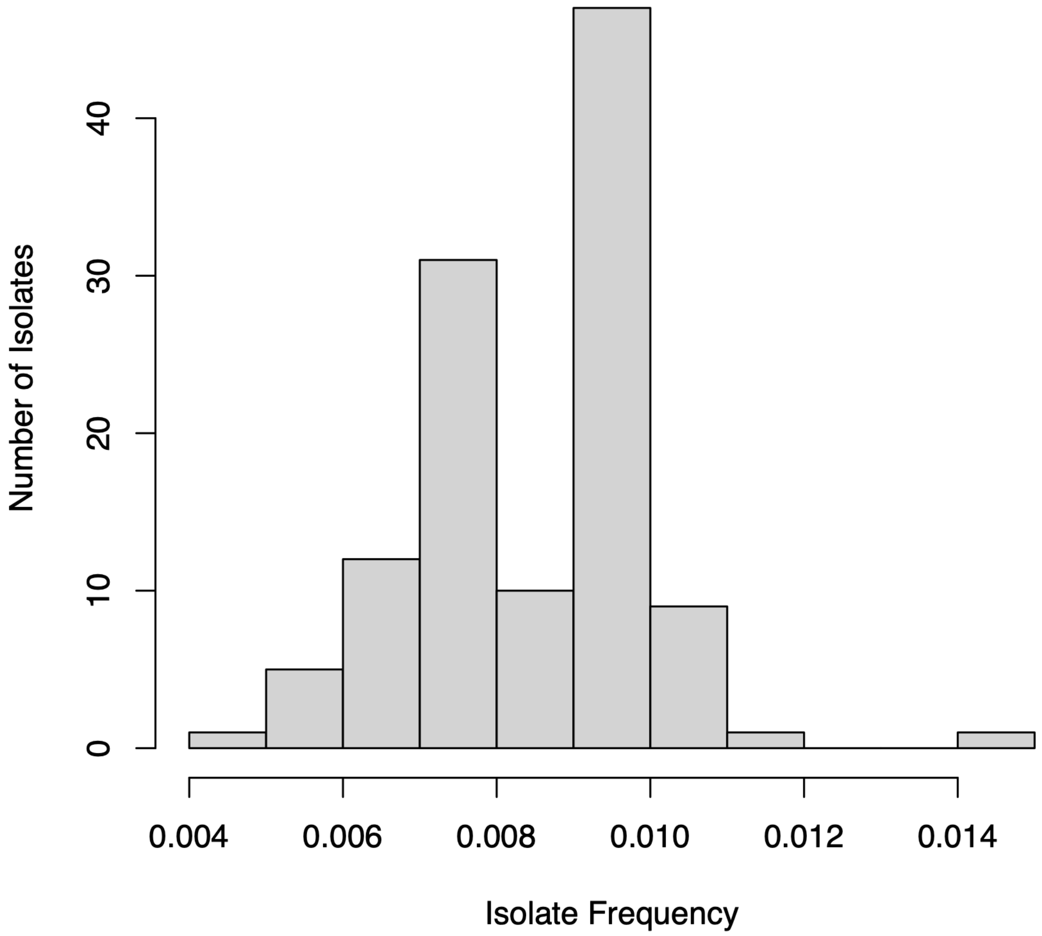


**Figure S5:** Histogram showing the distribution of the frequencies of 117 *S. meliloti* strains in the initial community used to inoculate the three alfalfa hosts.

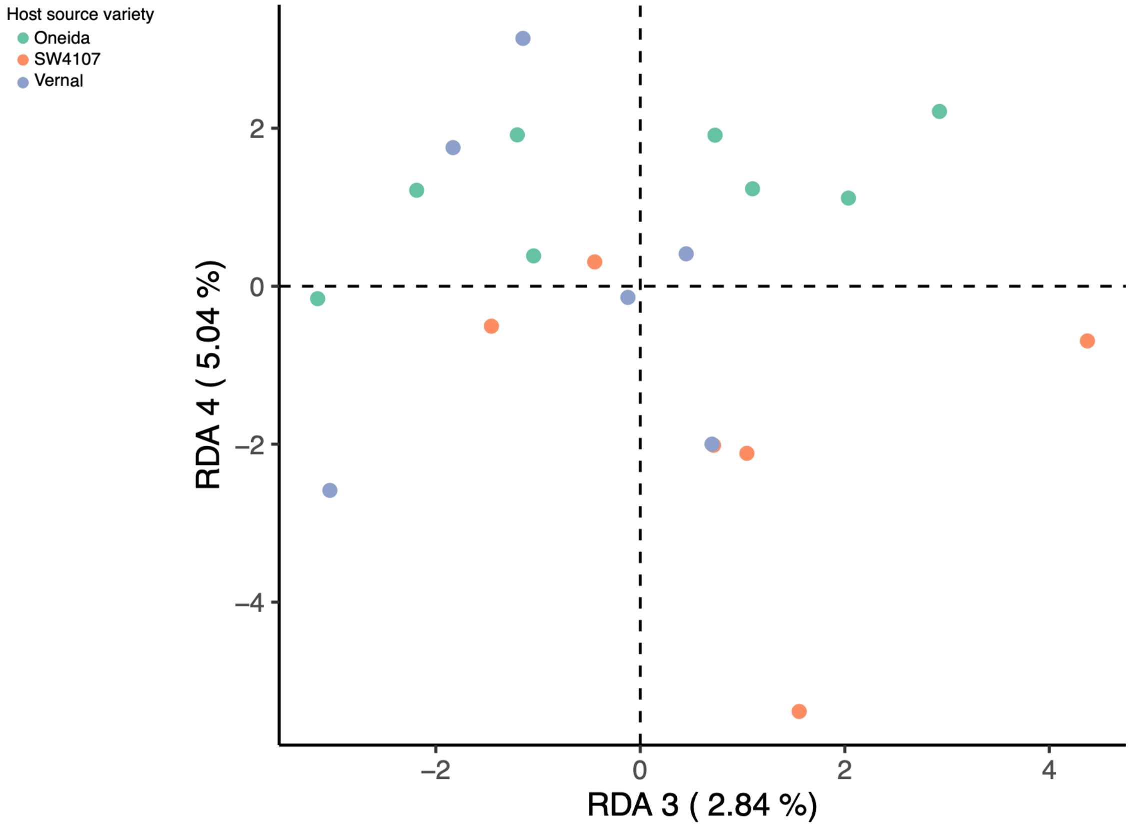


**Figure S6:** Redundancy Analysis (relative strain fitness ~ host) showing the third and fourth RDA axes, which explains 7.88% of the variation in the relative strain fitness. Individual points denote replicate pots, colours are coded based on the host variety.

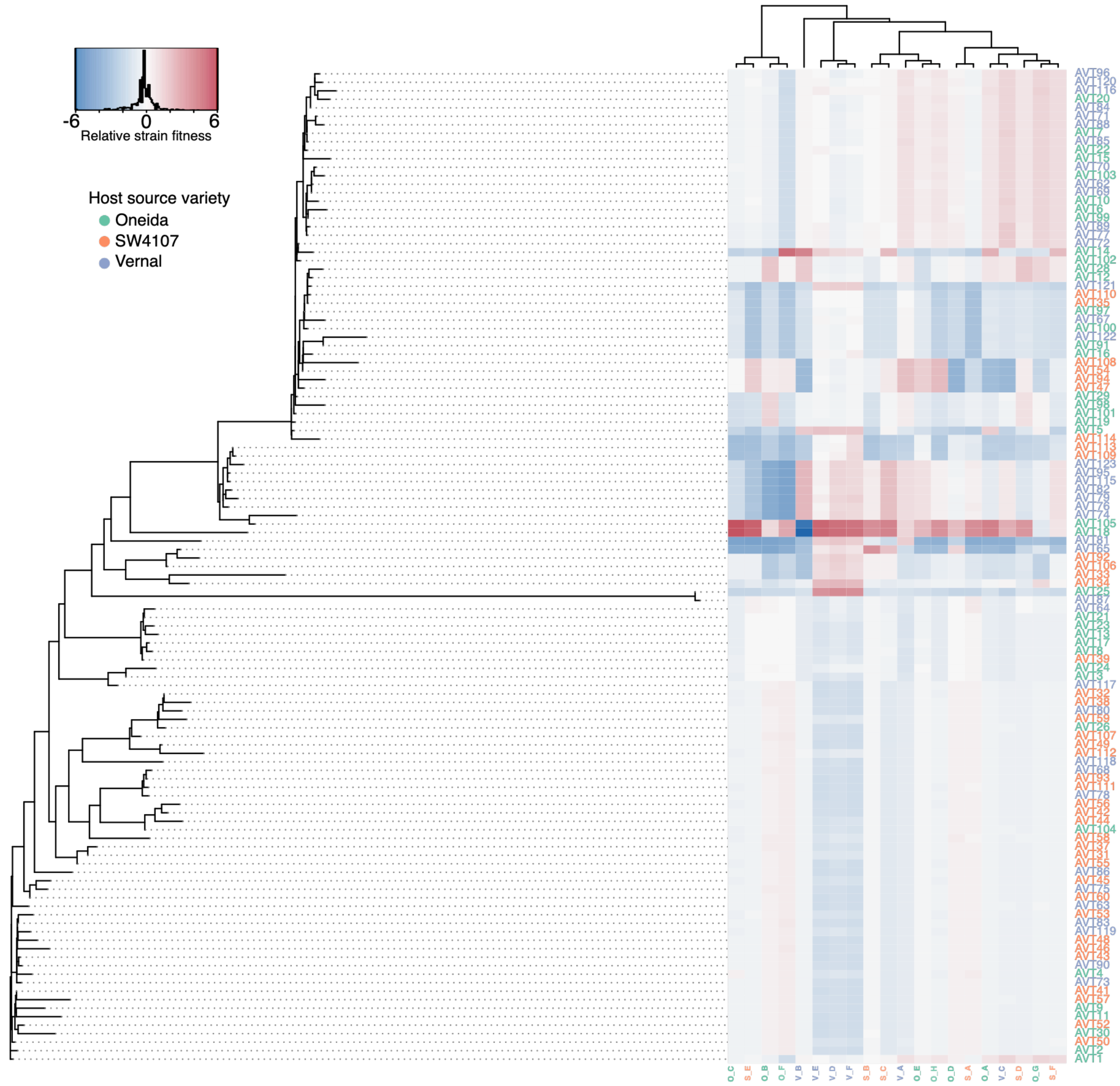


**Figure S7:** Heat map of relative strain fitness (strain log2-fold change) across the individual nodule pools from each plant (6 plants per variety), ordered according to the maximum-likelihood-based core gene phylogeny of the AVT strains (See Fig 2 for additional information). Strains enriched relative to their initial inoculum (Fig S4) appear in shades of red; depleted strains appear in shades of blue. The individual nodule pools are colour-coded based on their source host variety [orange=SW4107(S), green= Oneida, purple = Vernal]. The nodule pools are hierarchically clustered based on similarity in strain fitness profiles.

**Table S2**: Distribution of the core, softcore, shell, and cloud genes within the AVT strains across the replicons

| Replicon | PAV_class[*1] | n (number of genes | Proportion |
|---|---|---|---|
| Chromosome | Cloud (< 15%) | 2207 | 34.4 |
| Chromosome | Core (>99%) | 3096 | 48.36 |
| Chromosome | Shell (>= 15%) | 880 | 13.74 |
| Chromosome | Soft Core (>= 95%) | 218 | 3.4 |
| Plasmid_other | Cloud (< 15%) | 627 | 65.5 |
| Plasmid_other | Shell (>= 15%) | 329 | 34.4 |
| pSymA | Cloud (< 15%) | 681 | 25.14 |
| pSymA | Core (>99%) | 3 | 0.1 |
| pSymA | Shell (>= 15%) | 921 | 34.01 |
| pSymA | Soft Core (>= 95%) | 1103 | 40.73 |
| pSymB | Cloud (< 15%) | 422 | 20.38 |
| pSymB | Core (>99%) | 826 | 39.9 |
| pSymB | Shell (>= 15%) | 334 | 16.13 |
| pSymB | Soft Core (>= 95%) | 488 | 23.57 |

[*1] PAV_class = Presence-Absence Variation class. Definition of gene category: Core (>99%),Soft Core (95%-99%); Shell (15%-95%); Cloud (< 15%); based on(Gonzalez-Diaz et al., 2022).

**Table S3:** Distribution of SNPs across the three replicons within the AVT strain collection

| Replicon | SNPs[1] | Indels[2] | Multiallelics[3] |
|---|---|---|---|
| Chromosome | 57076 | 782 | 1586 |
| pSymA | 34349 | 586 | 279 |
| pSymB | 48722 | 1155 | 1159 |

[1]SNP= Single nucleotide polymorphism
[2]Indels= Insertions and deletions
[3]Multiallelic= Genomic loci with three or more observed alleles

**Table S4:** Normalized SNP (biallelic) counts across the replicons when the AVT strains were mapped to the in-house reference strain AVT118.

| Replicon | Median length | Raw SNP count | Normalized(SNPs/kb) |
|---|---|---|---|
| Chromosome | ~3.91 | 57076 | 14.58 SNPs/kb |
| pSymA | ~1.63 | 34349 | 18.24 SNPs/kb |
| pSymB | ~1.7 | 48722 | 29.44 SNPs/kb |

**Table S7:** Redundancy analyses of relative strain fitness followed by permutation-based ANOVA (Strain fitness ~ Host+Rep).

| Response | Pred[1] | %Exp[2] | Df[3] | Fval[4] | Pval[5] | R.adj[6] |
|---|---|---|---|---|---|---|
| Relative Strain fitness | Host | 36.17 | 2 | 3.35 | 0.01 | 0.11 |
| Relative Strain fitness | Rep[7] | 25.9 | 7 | 0.68 | 0.81 | NA |
| Relative Strain fitness | Residual | 53.87 | 10 | NA | NA | NA |

[1]Pred.= Predictors.
[2]%Exp = percentage of total variance explained by each predictor.
[3]Df = degrees of freedom.
[4]Fval = F statistic from the ANOVA model.
[5]Pval = significance level of the predictor effect.
[6]Radj = adjusted $R^2$ for the full model.
[7]Rep=Replicate
NA = not applicable.

**Table S8:** Pairwise RDA analysis and ANOVA-like permutation tests of relative strain fitness.

| Comparison | R.adj[1] | %Exp[2] | F.val[3] | P.val[4] | Host |
|---|---|---|---|---|---|
| O vs V | 0.19 | 0.34 | 6.2 | 0.002 | Significant* |
| O vs S | -0.0008 | 0.07 | 0.98 | 0.37 | None |
| V vs S | 0.12 | 0.20 | 3.7 | 0.01 | Significant* |

[1]Radj = adjusted $R^2$ for the full model
[2]%Exp = percentage of total variance explained by each predictor.
[3]Fval = F statistic from the ANOVA model.
[4]Pval = Significance level of the pairwise comparison.
*= statistically significant
